# Population structure and genetic diversity of the Critically Endangered bowmouth guitarfish (*Rhina ancylostomus*) in the Northwest Indian Ocean

**DOI:** 10.1101/2024.03.15.585225

**Authors:** Marja J. Kipperman, Rima W. Jabado, Alifa Bintha Haque, Daniel Fernando, P.A.D.L Anjani, Julia L.Y. Spaet, Emily Humble

## Abstract

The bowmouth guitarfish (*Rhina ancylostomus*) is a unique and understudied species of wedgefish with a distribution spanning the Indo-Pacific Oceans. Due to targeted and bycatch fisheries, this species is experiencing serious declines across its range. It is now considered among the most threatened species of elasmobranch. Despite this, species-specific management is limited, particularly around primary fishing hotspots. This is in part due to knowing very little about fundamental population processes. Here, we combine mitochondrial and single nucleotide polymorphism (SNP) data to carry out the first genetic assessment of *R. ancylostomus* across the Northwest Indian Ocean. We detect genetic differentiation across the northwest range, shaped by both oceanographic barriers and intrinsic dispersal constraints, and uncover a cline in genetic variation from east to west. These findings emphasise the importance of maintaining habitat connectivity while also identifying regions that may require heightened protection. In doing so, our study provides a critical baseline for conservation planning of *R. ancylostomus* and highlights the value of genomic data in elasmobranch conservation.

## Introduction

Characterising genetic diversity and population structure across extant species’ ranges is crucial for both *in situ* and *ex situ* conservation management (Hohenlohe et al., 2021). However, for many threatened species, such baseline information is severely lacking (Borgelt et al., 2022; Hochkirch et al., 2021; Tella et al., 2013). This compromises both the development and effectiveness of conservation actions. For example, the appropriate geographic scale of management largely depends on the extent to which populations are connected across space (Lowe & Allendorf, 2010; Mills & Allendorf, 1996; Mönkkönen & Reunanen, 1999). Discrete populations exchanging limited genetic information will benefit from local or regional scale management and may warrant delineation as conservation units (Moritz, 1999; Palsbøll et al., 2007). In contrast, populations connected by dispersal will require a more coordinated approach across larger spatial scales.

Large marine organisms exhibit contrasting patterns of genetic connectivity (Bailleul et al., 2018; Bernard et al., 2021, 2025; Pirog et al., 2019) Pelagic species with high dispersal capabilities frequently display high levels of gene flow across continuous distributions (Palumbi, 2003; Waples, 1998). This presents a challenge for conservation since it is not always clear where to delineate management units (Turbek et al., 2023). For example, individuals at opposite ends of a distribution may display distinct genetic and phenotypic variation yet remain connected by populations continuously distributed across space (Burri et al., 2016; Devitt et al., 2011; Irwin et al., 2005). Conversely, species exhibiting high site-fidelity and restricted movement are more likely to develop genetic differentiation between populations. If certain populations become small and isolated, they face increased vulnerability to exploitation, primarily due to heightened risks of inbreeding and loss of adaptive potential resulting from reduced genetic variation (Kimura et al., 1963; Lande, 1993; Lande & Shannon, 1996). From observational monitoring alone, it can be virtually impossible to identify such populations, especially in a marine environment.

Genetic and genomic tools provide a fantastic opportunity to explore the landscape of genetic diversity and differentiation (Bertola et al., 2024; Funk et al., 2012; Stapley et al., 2010) and are increasingly being applied to elusive marine megafauna (Kelley et al., 2016). For example, recent work has uncovered contrasting patterns of genetic diversity in two recently diverged manta ray (Mobulidae) species (Humble et al., 2024), signals of inbreeding in great hammerhead sharks (*Sphyrna mokarran*) (Stanhope et al., 2023) and subtle stock structure in blue shark (*Prionace glauca*) – a species previously thought to comprise a single global population (Nikolic et al., 2023). These insights all have the potential to influence management through highlighting populations in need of prioritisation, contributing to non-detriment findings under the Convention on International Trade of Endangered Species of Wild Flora and Fauna (CITES) and informing fisheries management (Casey et al., 2016; Domingues et al., 2018).

Guitarfish and wedgefish are among the most vulnerable and understudied groups of elasmobranchs (Dulvy et al., 2014; Kyne et al., 2020; Moore, 2017; Pytka et al., 2023). Their mostly coastal and in-shore habitat overlaps with much of the world’s most intensive fishing activities making them highly susceptible to bycatch (Jabado, 2018; White et al., 2013; Pytka et al. 2023), while international demand for their sought-after fins and thorns drive targeted exploitation (Clarke et al., 2006; Pytka et al., 2023). Consequently, ‘shark-like rays’ have been listed on Appendix II of CITES which seeks to regulate commercial trade, including of their derivative products. Among the most valuable fins on the market are those of the bowmouth guitarfish (*Rhina ancylostomus*), an evolutionarily distinct wedgefish (Gumbs et al., 2024) with the most widespread and continuous distribution of all shark-like rays (Figure 1). This species is listed as Critically Endangered on the IUCN Red List of Threatened Species and is particularly susceptible to population decline due to its slow growth, late maturity, and low fecundity (Kyne et al., 2019). Despite this, *R. ancylostomus* receives little to no species-specific management across its entire range, putting the species at serious risk of extinction.

**Figure 1.**
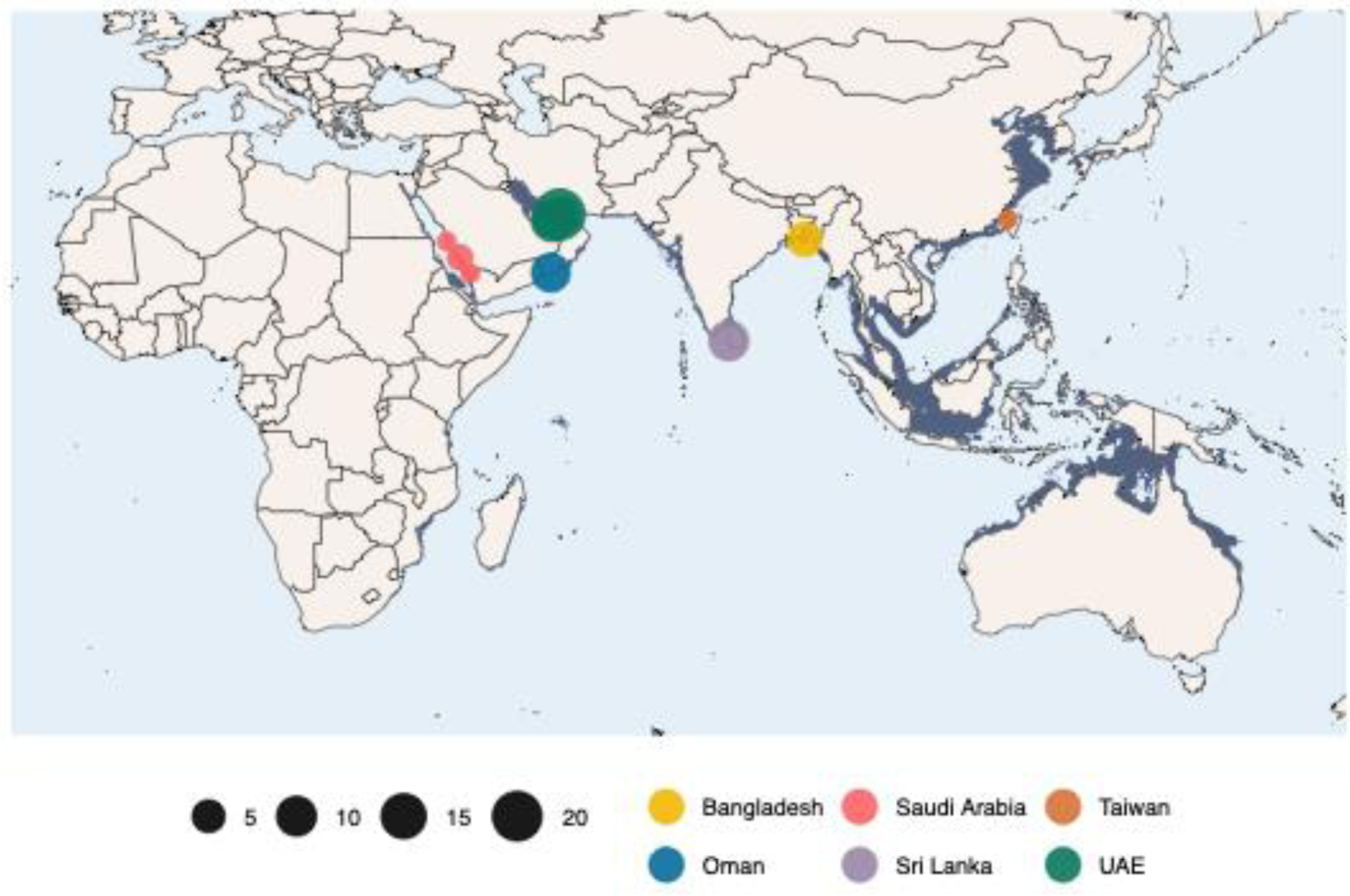
Geographic distribution of *R. ancylostomus* (dark blue) visualised together with the locations of samples used in this study. Sampling location points are distinguished by colour and scaled by the number of samples. The sample from Taiwan represents the *R. ancylostomus* mitochondrial genome assembly downloaded from NCBI and included in our mitochondrial DNA analysis. Further details are provided in Table S1.

Here, we carry out the first genetic assessment of *R. ancylostomus* across the Northwest Indian Ocean. We use a combination of mitochondrial and single nucleotide polymorphism (SNP) markers to build an atlas of genetic variation in order to address the following questions: (i) to what extent are populations genetically connected across the distribution; (ii) what are the levels of genetic variation at both mitochondrial DNA and SNPs and how is this partitioned among populations; and (iii) what are the implications of our findings for conservation management.

## Materials and Methods

### Sampling and DNA extraction

Fin clips were collected from 70 *R. ancylostomus* individuals from five geographic locations across the Northwest Indian Ocean: Saudi Arabia (*n* = 4), United Arab Emirates, Persian Gulf (*n* = 40), Oman (*n* = 12), Sri Lanka (*n* = 9) and Bangladesh (*n* = 5) (Figure 1). In all cases, samples were obtained from specimens captured in fisheries and stored in 99% ethanol at −22°C. Total genomic DNA was extracted using the Qiagen DNeasy® Blood and Tissue Kit following manufacturer protocols and quantified using both a Qubit dsDNA Broad Range Assay and running on a 1% agarose gel.

### Mitochondrial sequencing

Two regions of the mitochondrial genome were targeted for sequencing: cytochrome oxidase I (COI) and the control region (CR). A 650 bp region of the COI was amplified using universal primers FishF2 (5’ TCGACTAATCATAAAGATATCGGCAC 3’) and FishR2 (5’ ACTTCAGGGTGACCGAAGAATCAGAA 3’) (Ward et al., 2005). PCRs were carried out using 10 μl DreamTaq Green PCR Master Mix (2X) (ThermoFisher Scientific), 6 μl nuclease free water, 1 μl of each primer and 2 μl of template DNA. The following PCR profile was used for amplification: initial denaturation of 3 min at 95°C; 35 cycles of 30 sec at 95°C, 30 sec at 55°C, and 30 sec at 72°C, followed by a final extension of 5 min at 72°C. To amplify the CR, we designed a new set of primers (RACR_F: 5’ TGGGCTGGCGAGAAATAACC 3’, RACR_R: 5’ TTTTCGTTTCGACCCAGGGG 3’) in Geneious Prime v11.0.14.1 using the *R. ancylostomus* mitogenome as a reference (GenBank accession number KU721837.1). PCRs were carried out using the same volumes as above, together with the following amplification profile: initial denaturation of 1 min at 95°C; 35 cycles of 30 sec at 95°C, 30 sec at 58°C, and 30 sec at 70°C, followed by a final extension of 5 min at 70°C. The target product size for the CR fragment was 560 bp. PCR products were run on a 1.5% agarose gel to determine if amplification was successful. Successful amplicons were purified using the Exo-CIP Rapid PCR Cleanup Kit (New England BioLabs®) and sent for bi-directional Sanger sequencing with Eurofins Genomics.

Sequences were manually checked and edited using Geneious v1.2.3. Forward and reverse reads for each individual and each mitochondrial DNA (mtDNA) target region were *de novo* assembled into consensus sequences. Prior to further processing, each COI and CR consensus sequence was blasted to the NCBI non-redundant reference database using megablast and an E-value threshold of 0.05 for species identification purposes. Any sequence whose top BLAST hit based on sequence similarity and bit score was not *R. ancylostomus* was removed from the dataset. Remaining COI and CR consensus sequences were then multiple aligned using the Clustal Omega algorithm in Geneious and trimmed to the same length for downstream analysis. To increase the geographic scope of our analysis, we added target regions from the *R. ancylostomus* mitochondrial reference genome to each alignment. This reference sequence was generated using a sample originating from the Taiwan Strait (Si et al., 2016).

### SNP genotyping and filtering

SNP genotyping was carried out by Diversity Arrays Technology (DArT) who implemented a reduced representation sequencing protocol (DArTseq) followed by SNP calling with the DARTsoft14 pipeline (Cruz et al., 2013; Kilian et al., 2012). Briefly, DNA from 65 of the samples identified as *R. ancylostomus* was digested using PstI and SphI enzymes followed by adaptor ligation for each individual. Enzymes were chosen by DArT for their ability to isolate highly informative, low copy fragments of the genome. Following competitive PCR amplification, the resulting library was 250 bp paired-end sequenced on an Illumina HiSeq2500. To obtain a high quality dataset for downstream analysis, we filtered the resulting SNP genotypes using the R package *dartR* (Gruber et al., 2018; Mijangos et al., 2022). For this, we removed SNPs with low reproducibility scores based on technical replicates, retained one SNP per locus to account for potential linkage in the dataset and removed SNPs with a read depth less than 5 or greater than 50, and with a genotyping rate less than 80%. We then removed individuals with a call rate below 96%, that displayed excess heterozygosity or that displayed high relatedness with any other individual in the dataset. To determine relatedness, we estimated KING, R0 and R1 coefficients (Waples et al., 2019) using NgsRelate v2 (Korneliussen & Moltke, 2015). These statistics are based on genome-wide patterns of identity by state sharing between two individuals. This analysis was carried out using a highly informative dataset whereby SNPs that deviated significantly from HWE with a p-value threshold of 0.001, with a minor allele frequency < 0.1 or a genotyping rate <0.9 had been removed using PLINK v1.90. The individual with the lowest genotyping rate from any pairing exhibiting high relatedness was removed from the final dataset (Figure S1). At this point, we applied a final filter to the data to remove any SNPs with a minor allele frequency (MAF) <0.03, equivalent to a minor allele count of 3.

### Population structure

To investigate population structure using mtDNA, we generated median joining haplotype networks with PopART v1.7 (Leigh & Bryant, 2015) based on both the COI and CR alignments. Haplotype accumulation curves (HACs) were then calculated to determine how well within-species diversity was captured with our sampling effort. For this, we used the R package *HACSim* to simulate the number of samples required to observe the full range of haplotype variation that exists for a species (Phillips et al., 2020). The simulation was run for 10,000 permutations, a confidence level of 0.95 and at different percentage recovery rates until no additional individuals were necessary for recovery of haplotypes (0.95). We further explored population differentiation by calculating pairwise *F*_ST_ between sampling locations using Arlequin v3.5.22 (Excoffier & Lischer, 2010). This was run separately for each mtDNA gene with 1000 permutations to estimate 95% confidence intervals. Finally, to measure population genetic structure both within and between groups, we ran an Analysis of Molecular Variance (AMOVA) using Arlequin and tested significance using 1000 permutations.

To investigate population structure using the SNP data, we implemented three approaches. First, we performed a principal components analysis using the R package *adegenet* (Jombart, 2008). Second, we used the Bayesian clustering algorithm in STRUCTURE to identify the number of genetic clusters (*K*) in the dataset. STRUCTURE was run for values of *K* ranging from *K* = 1 – 8, with ten simulations for each *K* and a burn-in of 50,000 iterations followed by 100,000 MCMC iterations. We used the admixture and correlated allele frequency models without sampling location information. We then used the R package pophelper (Francis 2017) to analyse the results, parse the output to CLUMPP for averaging across simulations and to visualise assignment probabilities. The optimal *K* was identified based on the maximum value of the mean estimated *ln* probability of the data (Ln Pr(*X* | *K*)) (Pritchard et al 2000) and the Δ*K* method (Evanno et al 2005). Finally, we estimated pairwise genetic differentiation between populations using the Weir and Cockerham *F*_ST_ value (Weir & Cockerham, 1984) in the R package dartR. Confidence intervals and *p-*values were estimated based on bootstrap resampling of loci within each sampling location 1000 times (Weir & Cockerham, 1984). To account for the effects of unbalanced sampling on our inference of population structure (Puechmaille, 2016), we randomly downsampled the UAE population to eight individuals, resulting in a more uniform distribution of samples across locations. We recalculated MAF and removed SNPs with a MAF <0.03 before re-running PCA, STRUCTURE and *F*_ST_ as described above. Results displayed similar patterns for the subsampled dataset as the full dataset (Figures S7–9) and therefore those for the full dataset are presented below.

### Isolation by distance

To investigate patterns of isolation by distance in *R. ancylostomus*, we explored the relationship between genetic and geographic distance between all pairs of populations. Geographic location was assigned to each individual by determining the coordinates of the coastline directly adjacent to the fisheries landing site in which the sample was collected. Although this is not the true location of where the individual was caught, it represents a fair approximation given the coastal nature of the species. Genetic distances were based on COI-based pairwise *F*_ST_ estimates as calculated above. We determined geographic distances based on a least-cost path analysis using the R package *marmap* (Pante & Simon-Bouhet, 2013). A minimum depth constraint of - 10 metres was applied to ensure no paths were overland. Significance between genetic and geographic distance matrices was then calculated using distance-based Moran’s eigenvector maps (dbMEM) by redundancy analysis (RDA, Legendre et al., 2015). Geographical distances were transformed into dbMEMs using the R package *adespatial*, and genetic distances were decomposed into principal components using the R function *prcomp*. RDA was then performed using the R package *vegan*, with significance tested using 1000 permutations.

### Genetic diversity and heterozygosity

Genetic diversity estimates were calculated for the mtDNA data using the software DnaSNP v6.12.03 (Rozas et al., 2017) at both a species and sampling location level. For the SNP data, we assessed levels of genetic variation by estimating multi-locus heterozygosity for each individual using the R package *inbreedR* (Stoffel et al., 2016).

## Results

### Mitochondrial sequencing

Out of a total of 70 samples, 66 were confirmed in a BLAST search using COI and / or CR sequences to originate from *R. ancylostomus*. The remaining four samples were identified as tiger shark (*Galeocerdo cuvier)*, painted sweetlips (*Diagramma pictum)*, starry puffer (*Arothron stellatus)* and *Pseudomonas campi* and were removed from further analysis. These results are likely due to mislabelling of samples. Of the 66 confirmed *R. ancylostomus* samples, a total of 63 successfully amplified at COI, and 65 successfully amplified at the CR. Final filtered alignment lengths were 674 bp for COI and 512 bp for the CR. Analysis of mitochondrial haplotypes revealed a total of eleven unique sequences for COI and three for the CR, including the sequence originating from the mitochondrial reference genome. Of these, ten and two haplotypes were newly discovered, respectively (Table 1). All measures of genetic diversity were consistently higher for COI compared to the CR (Table 1).

**Table 1.** Genetic diversity measures for *R. ancylostomus* calculated from COI and CR sequences generated as part of this study.

| Indices | COI | CR |
| --- | --- | --- |
| Number of sequences | 63 | 65 |
| Number of segregating sites (*S*) | 18 | 4 |
| Number of haplotypes (*H*) | 10 | 3 |
| Haplotype diversity (*Hd*) | 0.65028 | 0.49135 |
| Nucleotide diversity (*pi*) | 0.00377 | 0.00181 |

### SNP genotyping

A total of 19,436 SNPs were genotyped in 65 individuals using the DARTsoft14 pipeline. After quality filtering, the final dataset contained 3,533 SNPs in 49 individuals from across all sampling locations (Saudi Arabia = 2, Oman = 8, UAE = 33, Sri Lanka = 5, Bangladesh = 1). For a full breakdown of the number of SNPs and individuals removed at each filtering step, see Table S2.

### Population structure

Haplotype networks for both gene regions revealed a noticeable geographic signal from west to east of the sampling range (Fig 2A–B). This pattern was more distinct for COI, where a greater number of haplotypes were observed overall. In particular, we observed no haplotype sharing between locations at the extremes of the sampling range while Sri Lanka, which was intermediate, shared haplotypes with every location except for Taiwan. We also observed geographic signal among samples collected in Sri Lanka, where individuals originating from the west coast shared no haplotypes with those originating from the east coast.

**Figure 2.**
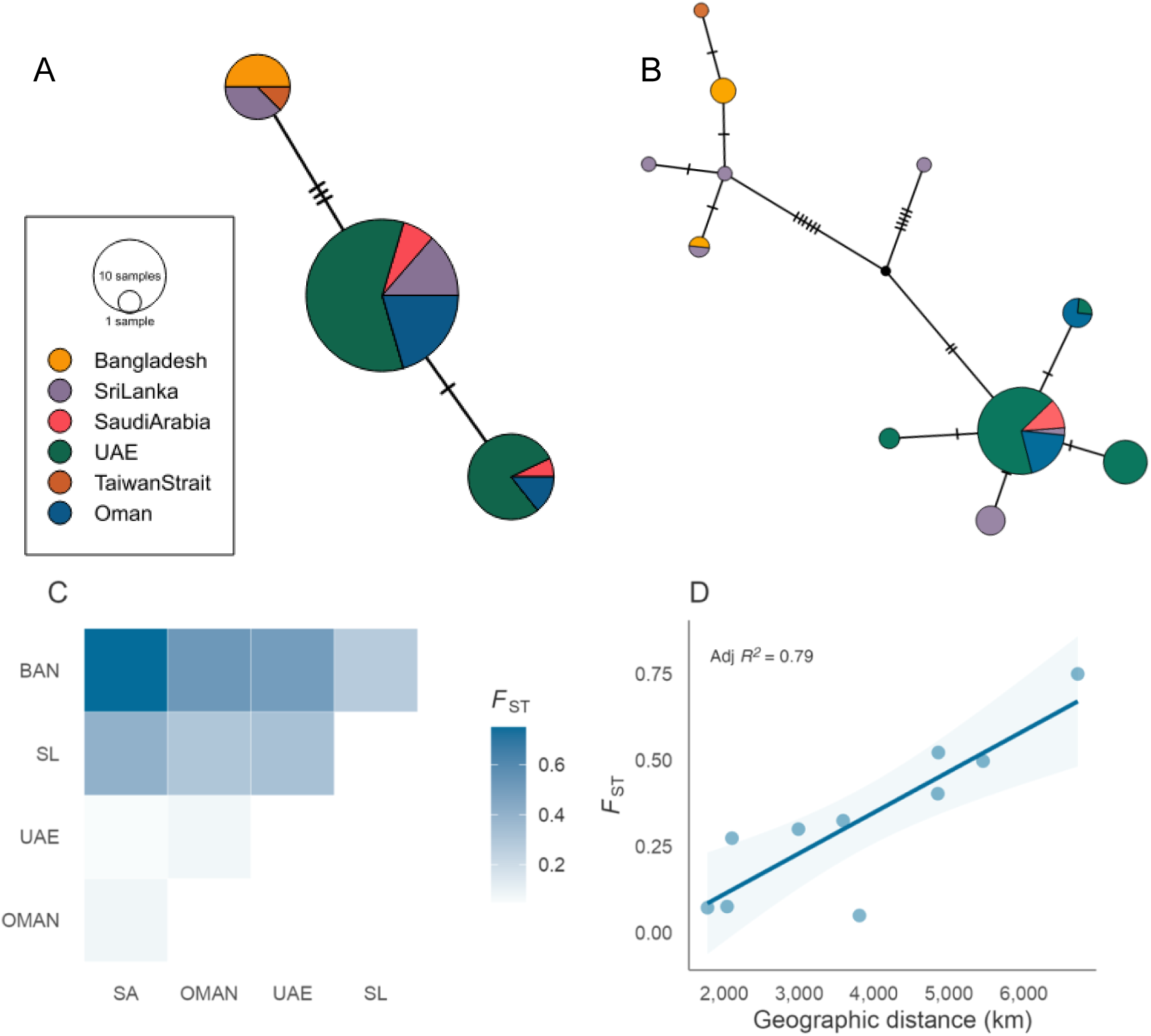
Mitochondrial-based population differentiation and isolation by distance in *R. ancylostomus*. Median-joining mitochondrial haplotype network for (A) CR and (B) COI sequences. Colours reflect geographic origin, node size is proportional to the number of individuals displaying each haplotype and hashed lines indicate the number of nucleotide substitutions between haplotypes. (C) Pairwise *F*_ST_ estimates between sampling locations based on COI. Population abbreviations: BAN, Bangladesh; SL, Sri Lanka; UAE, United Arab Emirates; OMAN, Oman; SA, Saudi Arabia. (D) Relationship between genetic (COI based *F*_ST_) and geographic distance as calculated by least-cost path analysis for all pairwise population comparisons. Solid lines and shaded areas reflect the regression slopes and standard errors, respectively, based on a linear model.

Based on haplotype accumulation curves (Figure S2), our COI dataset represents around 85% of the haplotype diversity in the species. An additional 63 and 132 individuals would be required to recover 95% and 99% of the haplotype diversity respectively (Figure S2A–B). For the CR, our sample size of 65 individuals was sufficient to recover both 95% and 99% of the haplotype diversity in the species (Figure S2C–D).

### Population differentiation

To further explore levels of differentiation, we carried out an AMOVA and calculated pairwise *F*_ST_ using both mitochondrial gene regions. AMOVA revealed that most of the observed variation is partitioned within populations for both COI and the CR (Table 2). However, there was also significant evidence of variation at a population level, with 29.07% and 25.58% of the variation occurring between populations for COI and CR respectively. Pairwise *F*_ST_ estimates ranged from 0.04–0.75 for COI and from −0.18–0.76 for the CR (Figure 2C and Table S3).

**Table 2.**
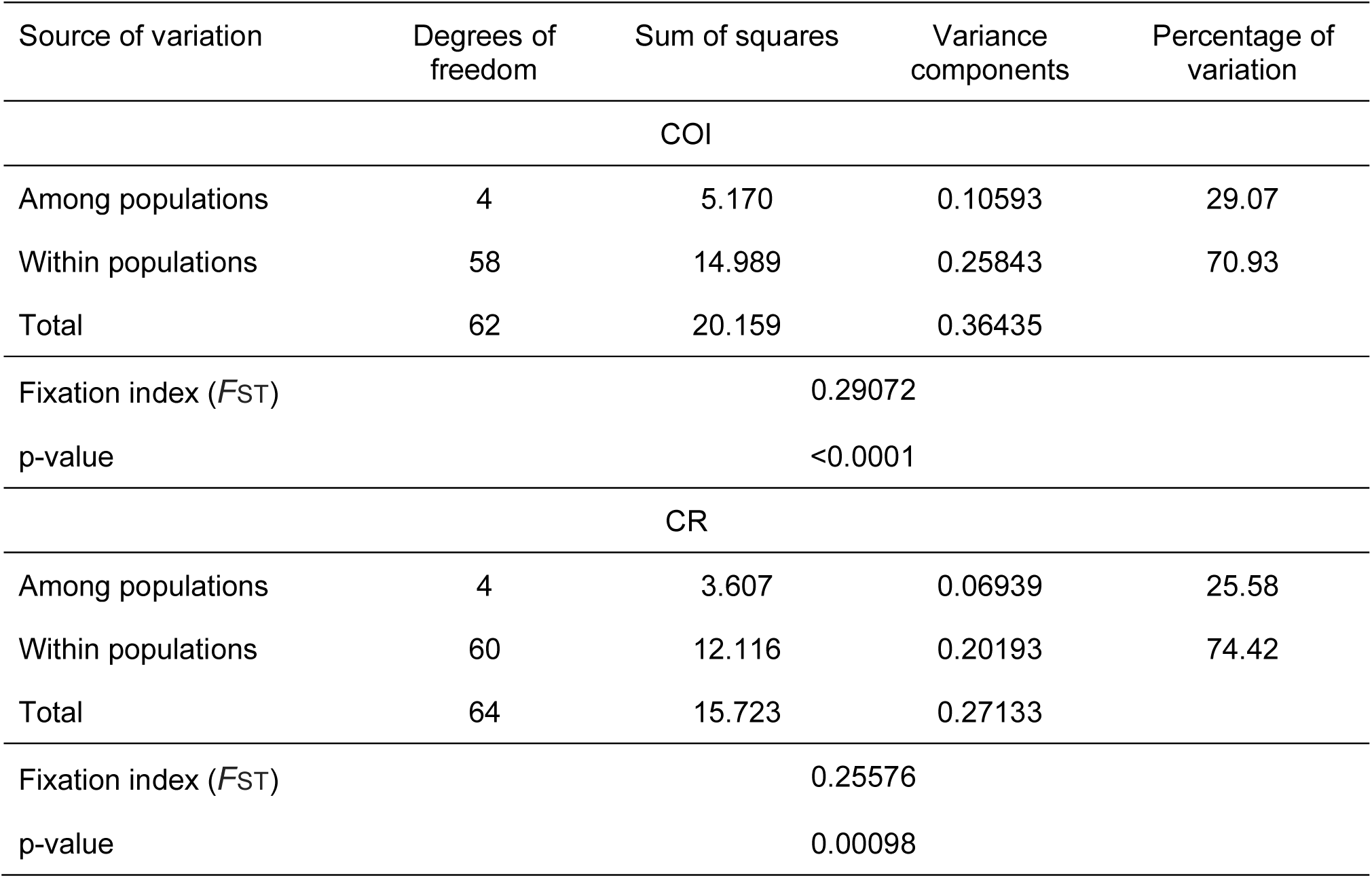
Summary of analysis of molecular variance (AMOVA) for COI and CR sequences in *R. ancylostomus*. Variation is partitioned among and within populations and a p-value is significant if <0.05.

Significant differences were observed between Bangladesh and all other populations for both COI and CR (Table S3). This was also the case for Sri Lanka at COI. In contrast, no comparisons between populations in the Arabian Peninsula were found to be significant suggesting an effect of geographic proximity. In line with this, although no significant relationship was observed between pairwise *F*_ST_ and geographic distance (adjusted *R2* = 0.79, *P* = 0.075) there was a clear tendency for populations separated by greater distances to display higher genetic differentiation indicating a signal of isolation by distance (Figure 2D).

### SNP results

Analysis of SNP data revealed a similar yet more pronounced pattern of population structure across the sampling range. In the PCA analysis, samples from most sites clustered apart along PC1 (Figure 3A). In particular, individuals from Oman and Saudi Arabia clustered separately from those originating from the UAE (Persian Gulf), with a deeper split observed between samples originating from the Red Sea, Arabian Sea and the Persian Gulf, and those originating from Sri Lanka and Bangladesh. No clustering was observed between sampling locations along PC2 (Figure 3A) or PC3 (Figure S3). These patterns were in part mirrored by the STRUCTURE analysis in which an optimal value of *K* = 2 was inferred based on Δ*K* and log likelihood values (Figure 3B, Figure S4–5). Clusters correspond to the east and west proximities of the sampling range with all individuals from Oman and Saudi Arabia being assigned to Cluster I with q > 0.85, and individuals from Sri Lanka and Bangladesh being assigned to Cluster II with q > 0.81.

**Figure 3.**
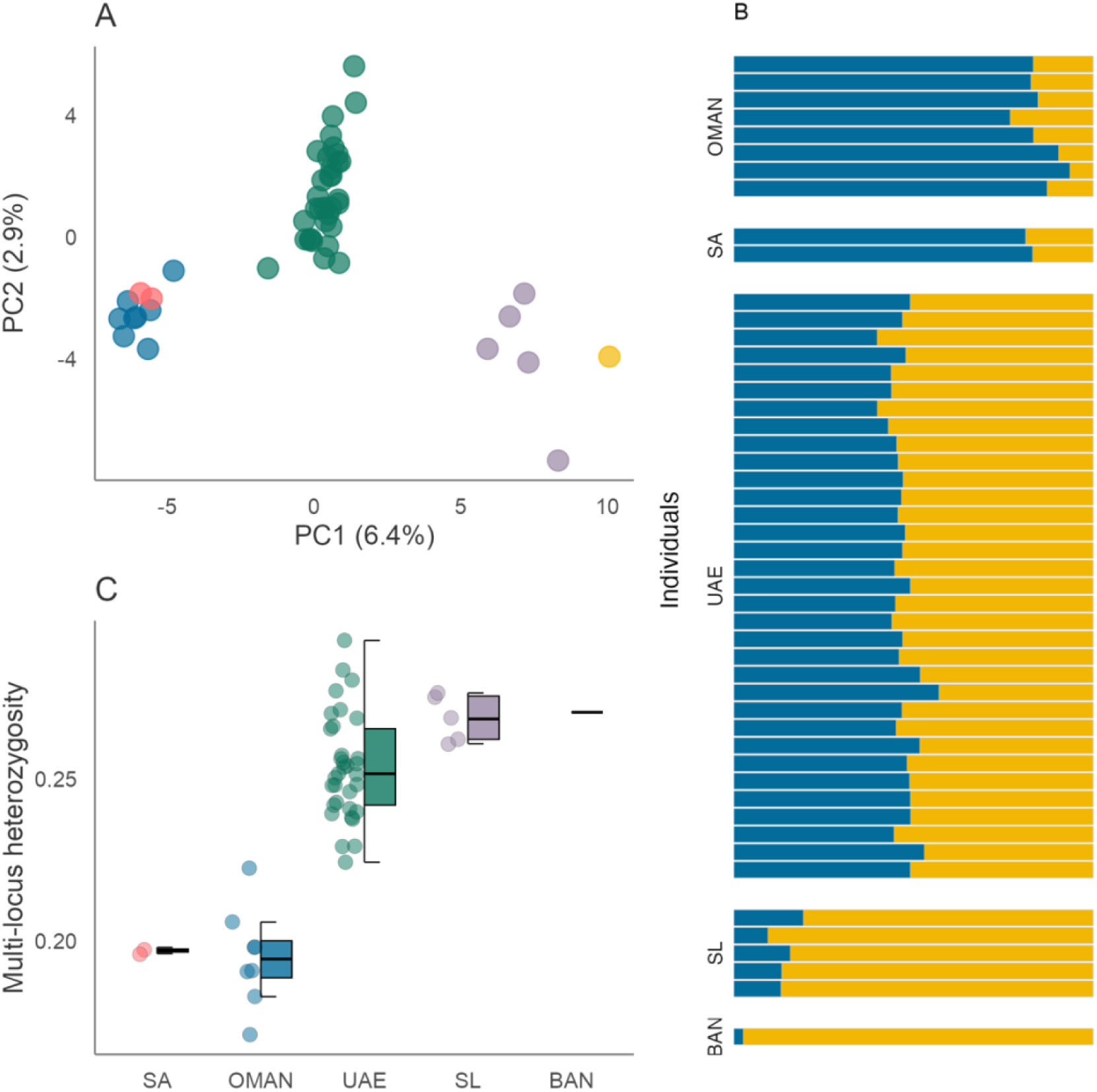
SNP-based population structure and diversity in *R. ancylostomus*. (A) Scatterplot showing individual variation in principal components (PC) one and two derived from principal components analysis. The amount of variance explained by each PC is shown in parentheses. Blue circles correspond to individuals from Oman, pink from Saudi Arabia, green from the UAE, purple from Sri Lanka and yellow from Bangladesh (B) Individual assignment to genetic clusters based on STRUCTURE analysis for *K* = 2 using 3,565 SNPs and 49 individuals. Each horizontal bar represents a different individual and the relative proportion of the different colours indicate the probabilities of belonging to each cluster. Individuals are separated by sampling locations as indicated in Figure 1. (C) Variation in multi-locus heterozygosity among populations. Centre lines of boxplots reflect the median, bounds of the boxes extend from the first to the third quartiles, and upper and lower whiskers reflect the largest and smallest values but no further than 1.5 x the interquartile range from the hinge. Population abbreviations: BAN, Bangladesh; SL, Sri Lanka; UAE, United Arab Emirates; OMAN, Oman; SA, Saudi Arabia

Individuals from intermediate locations in the UAE displayed admixture between inferred clusters (Cluster I mean q = 0.53; Cluster II mean q = 0.47) suggesting some degree of gene flow across the range. Pairwise *F*_ST_ estimates ranged from 0.01–0.27 and tended to be higher between populations separated by greater distances, in line with our mitochondrial analysis (Figure S6). All estimates including Bangladesh had larger confidence intervals than those without, most likely due to the small sample size of this population.

### Genetic diversity

To explore the landscape of genetic variation, we calculated genetic diversity estimates across sampling locations. Haplotype and nucleotide diversity at COI were highest for individuals originating from Sri Lanka and Bangladesh (Table 4). Individuals from the UAE and Oman had intermediate levels of diversity, while the lowest levels of diversity were observed in Saudi Arabia where only one haplotype was detected. Similar patterns were observed at the CR except that Bangladesh displayed the lowest variation and Saudi Arabia was intermediate.

**Table 4.**
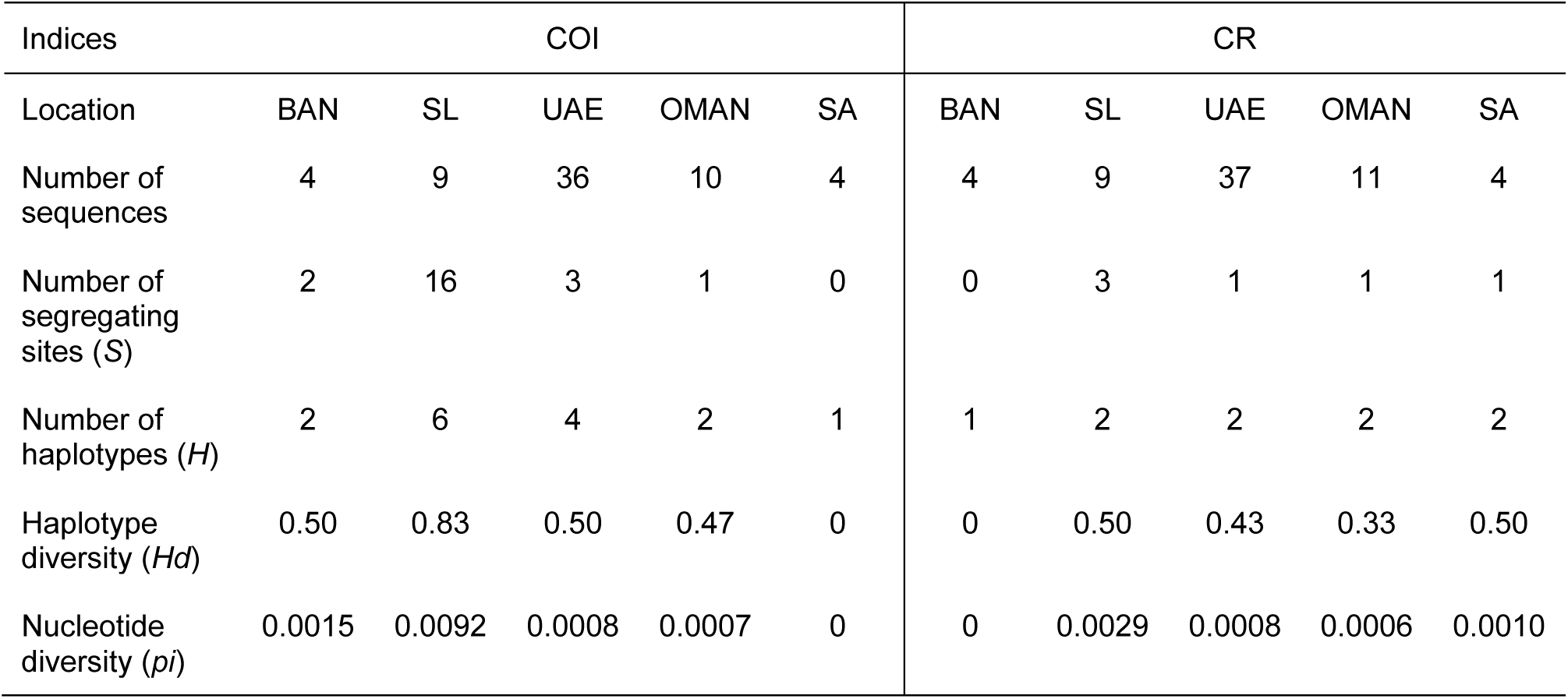
Genetic diversity measures for each *R. ancylostomus* sampling location calculated from COI and CR sequences.

| Indices | COI |  |  |  |  | CR |  |  |  |  |
| --- | --- | --- | --- | --- | --- | --- | --- | --- | --- | --- |
| Location | BAN | SL | UAE | OMAN | SA | BAN | SL | UAE | OMAN | SA |
| Number of sequences | 4 | 9 | 36 | 10 | 4 | 4 | 9 | 37 | 11 | 4 |
| Number of segregating sites (*S*) | 2 | 16 | 3 | 1 | 0 | 0 | 3 | 1 | 1 | 1 |
| Number of haplotypes (*H*) | 2 | 6 | 4 | 2 | 1 | 1 | 2 | 2 | 2 | 2 |
| Haplotype diversity (*H<sub>d</sub>*) | 0.50 | 0.83 | 0.50 | 0.47 | 0 | 0 | 0.50 | 0.43 | 0.33 | 0.50 |
| Nucleotide diversity (*p<sub>i</sub>*) | 0.0015 | 0.0092 | 0.0008 | 0.0007 | 0 | 0 | 0.0029 | 0.0008 | 0.0006 | 0.0010 |

When comparing individual heterozygosity calculated from SNP data, we detected a noticeable decrease in variation from east to west of the sampling range (Figure 3C). In line with findings from the mtDNA analysis, individuals originating from Saudi Arabia and Oman displayed the lowest levels of variation, with little overlap in heterozygosity values between any of the other populations. Individuals from Sri Lanka and Bangladesh displayed the highest levels of variation. Unsurprisingly, the greatest variance in heterozygosity was observed in individuals originating from the UAE, likely driven by its high sample number.

## Discussion

*Rhina ancylostomus* are among the most threatened elasmobranch species. While some international measures are in place to reduce pressure on this species, national and regional management is hindered in part due to gaps in our understanding of population connectivity and genetic variation. We used a combination of mitochondrial and SNP data to shed light on population processes in and around the Northwest Indian Ocean. Our findings have implications for both *in situ* and *ex situ* management and conservation planning and provide a solid baseline with which to build our knowledge of this threatened and understudied species.

Our results first highlighted an unexpected difference in haplotype variation between COI and the CR. Across many vertebrates, the CR is the most polymorphic region of the mitochondrial genome (McMillan & Palumbi, 1997; Saccone et al., 1991; Zhang et al., 1995) making it a common marker for investigating intraspecific variation (Avise, 1995). It was therefore somewhat surprising to observe lower levels of variation at the CR than at COI across multiple *R. ancylostomus* populations. However, in many teleost fish it is well established that the CR evolves at a similar rate to the whole mitochondrial genome (Bernatchez & Danzmann, 1993; Shedlock et al., 1992; Zhu et al., 1994), and similar patterns have more recently been observed in other elasmobranch species (Boomer et al., 2012; Castillo-Páez et al., 2014; Dudgeon et al., 2009). While it is possible we inadvertently targeted a conserved sequence domain, our findings indicate this pattern could extend into numerous shark and ray families. A possible driver of low CR variation includes functional constraints for a particular base composition and therefore selection against transition mutations (Apostolidis et al., 1997; Bernatchez & Danzmann, 1993). Further work comparing complete gene regions and whole mitogenomes across species and populations will shed light on the extent and mechanisms of mitochondrial evolutionary rate variation in elasmobranchs. Nevertheless, those interested in intraspecific variation in related species will benefit from targeting highly variable regions of the mitochondrial genome such as COI and NADH2 (Naylor et al., 2012).

To explore patterns of intraspecific variation, we supplemented our mtDNA with SNP genotypes. Both marker sets highlighted some degree of genetic variation across the Northwest Indian Ocean, largely between individuals from Saudi Arabia, Oman and the UAE, and those from Sri Lanka, Bangladesh and Taiwan. The Indian Ocean Barrier is an established barrier to gene flow, particularly for coastal elasmobranchs (Hirschfeld et al., 2021), and likely contributes to the large-scale differences observed here. At the same time, SNP-based structure and PCA analysis suggested finer-scale clustering associated with sampling location. Such patterns may also reflect the species’ intrinsic dispersal limitations, with restricted movement leading to gradual shifts in allele frequencies across the range. However, it is important to acknowledge that sample acquisition from this endangered species is challenging, resulting in limitations to our sampling design. In addition to the locations sampled as part of this study, *R. ancylostomus* occurs along the coastlines of all countries in between, including eastern Saudi Arabia, Iran, India, and Pakistan (Kyne et al., 2019, Figure 1). Uneven sampling in a continuously distributed species can lead to the presence of artificial clusters due to spatial autocorrelation in allele frequencies (Chambers & Hillis, 2020; Perez et al., 2018). Population structure observed in our dataset may therefore be driven by gaps in our sampling distribution as opposed to true population boundaries. We therefore interpret the observed structure as arising from a combination of extrinsic barriers such as the Indian Ocean Barrier and intrinsic dispersal constraints, producing a pattern of isolation by distance across much of the range assessed here.

The distribution of *R. ancylostomus* extends beyond the Northwest Indian Ocean, down the coast of East Africa in one direction (including offshore islands such as the Seychelles and Reunion), and across to South East Asia, Australia and the East China Sea in the other (Kyne et al., 2019). While our study provides a significant advance in our understanding of the species, sampling constraints prevented assessment across its full range, limiting our ability to characterise the complete landscape of genetic variation. Building on the patterns observed in the Northwest Indian Ocean, two scenarios seem plausible across the broader distribution. First, decreasing genetic similarity with increasing genetic distance may extend across the entire range, consistent with the continuous nature of the species distribution and the intrinsic dispersal constraints that likely drive isolation by distance. There are a handful of examples demonstrating such patterns in coastal sharks, including the lemon shark (*Negaprion brevirostris*), sandbar shark (*Carcharhinus plumbeus)* and Caribbean reef shark (*Carcharhinus perezi,* Ashe et al., 2015; Bernard et al., 2017; Pember et al., 2023), although complete distributions are rarely assessed. Yet, even under such scenarios, adaptive loci may show signatures of differentiation despite clinal variation at neutral loci (Turbek et al., 2023). Second, extrinsic barriers to gene flow can lead to breakpoints in patterns of isolation by distance. For example, deep water trenches and strong ocean currents have been shown to restrict movements and limit gene flow in many coastal elasmobranchs (Dudgeon et al., 2009; Hirschfeld et al., 2021; Humble et al., 2024; Schultz et al., 2008). Given its propensity for shallow inshore environments, we anticipate barriers to gene flow beyond the Northwest Indian Ocean will likely give rise to population differentiation in *R. ancylostomus*. Nevertheless, one record exists of an individual caught by a purse seiner in pelagic waters between the African continent and the Seychelles (Forget & Muir, 2021) suggesting the species may be capable of longer distance movements. Further sampling and investigation across the entire species range, while challenging, will be crucial to disentangle the relative roles of dispersal limitation and oceanographic barriers in shaping population structure.

In addition to investigating population structure, we also explored the landscape of genetic variation among populations. Findings from SNP data, and to some extent mitochondrial haplotypes, highlighted a decline in variation from east to west of the sampling range. Such a pattern may have emerged as a result of species range expansion, a process which inherently impacts diversity as repeated founder events drive the loss of low frequency alleles (Le Corre & Kremer, 1998; Peter & Slatkin, 2013). While we cannot infer the true geographic origin of the species, we hypothesise this may have occurred somewhere in the Indo-Malayan archipelago. This region has extraordinary levels of biodiversity and is thought to act as a centre of origin and survival for many marine species (Evans et al., 2016), including some coast elasmobranchs (Maisano Delser et al., 2019; Walsh et al., 2022). With this in mind, we cautiously expect populations further east of the sampled range to harbour higher levels of genetic diversity and those along the east African coast to harbour lower levels. In line with this, Saudi Arabia and Oman displayed the lowest levels of individual variation in our study. Further sampling in these regions, particularly at range edges, will be crucial to validate our results and to explore this hypothesis in more detail. Genetic variation is fundamental not only for buffering the intrinsic impacts of population decline but for enabling populations to adapt to extrinsic pressures (Bonnet et al., 2022; Lai et al., 2019). Given the intensity of fishing pressure across the Arabian Sea and its adjacent waters (Jabado et al., 2018; Jabado & Spaet, 2017; Spaet & Berumen, 2015) our results paint a worrying picture for the future of the species in this region.

### Conservation implications and future directions

Guitarfish and wedgefish are notoriously neglected in conservation management (D’Alberto, 2022; Kyne et al., 2019; Pytka et al., 2023). While some local and national fisheries management measures can provide indirect benefits, there are few species-specific measures in place. By advancing our understanding of *R. ancylostomus* populations, our results provide fundamental information for guiding targeted management and conservation planning for the species. We uncover genetic differentiation across the Northwest Indian Ocean that is likely explained by both intrinsic dispersal constraints and extrinsic barriers to gene flow. Such processes are common in coastal marine taxa, producing gradual clines in allele frequencies in some areas but more distinct breaks where oceanographic barriers, such as the Indian Ocean

Barrier, restrict connectivity. These dynamics complicate the designation of discrete management units (Turbek et al., 2023) but highlight the importance of maintaining habitat connectivity across the species’ northern range, combined with local and national fisheries management. The former is crucial for maximising genetic diversity and buffering the impacts of local exploitation while the latter recognizes how fisheries management ultimately works in practice. We also identify regions with markedly lower genetic variation that should warrant heightened protection. Recently designated Important Shark and Ray Areas (ISRAs) in the Western Indian Ocean (Jabado et al., 2024) not only identify these locations but partly capture the continuous coastal distribution of *R. ancylostomus* and should therefore help kickstart the development of appropriate area-based conservation management.

Our findings also have implications for *ex situ* management. *Rhina ancylostomus* is one of few wedgefish species routinely held in aquarium collections across the globe (Smith et al., 2017). Through building an atlas of genetic variation for a key part of the species range, we provide a valuable reference for determining the breadth of variation present in the *ex situ* collection.

Due to the risks associated with small population size in captivity, such knowledge is crucial for breeding, movement and supplementation decisions (Frankham et al., 2002; Lacy, 1987). While there are excellent examples of well managed terrestrial collections (Farquharson et al., 2022; Humble et al., 2023; Witzenberger & Hochkirch, 2011), the use of genetic information in *ex situ* management of marine species significantly lags behind. *Rhina ancylostomus* therefore presents an excellent opportunity to integrate *ex situ* and *in situ* genetic management in an aquatic species under a One Plan approach to conservation (Pritchard et al., 2012; Redford et al., 2012; Schwartz et al., 2017). Nevertheless, we acknowledge the potential existence of sub-populations in areas of the species range not assessed here. If *R. ancylostomus* becomes a candidate for restoration or reintroduction, this information will be fundamental for establishing and managing an insurance population, identifying release sites and selecting individuals for release. We therefore strongly advocate for further sampling and investigation – particularly around East Africa and South East Asia– in order to support holistic and coordinated conservation management of this unique and threatened species.

## Acknowledgements

We thank all the volunteers that generously provided their time to support with data collection in the United Arab Emirates and Bangladesh. We are grateful to the Department of Wildlife Conservation and the Department of Fisheries and Aquatic Resources in Sri Lanka for their advice and support, and to all the BRT researchers and volunteers for their long hours at landing sites.

## Data, scripts, code and supplementary information availability

Data analysis code is available at https://github.com/mkp131/Bowmouth_analysis and https://github.com/elhumble/rhina_popgen_2024. Mitochondrial haplotypes have been deposited on NCBI (PV468213– PV468222; PV467736– PV467738). SNP genotypes and metadata are available at https://github.com/elhumble/rhina_popgen_2024.

