## Supplementary Material for "Population structure and genetic diversity of the Critically Endangered bowmouth guitarfish (*Rhina ancylostomus*) in the Northwest Indian Ocean"

### Table of contents

Supplementary Figures **Page 2**

Supplementary Tables **Page 10**

### Supplementary Figures

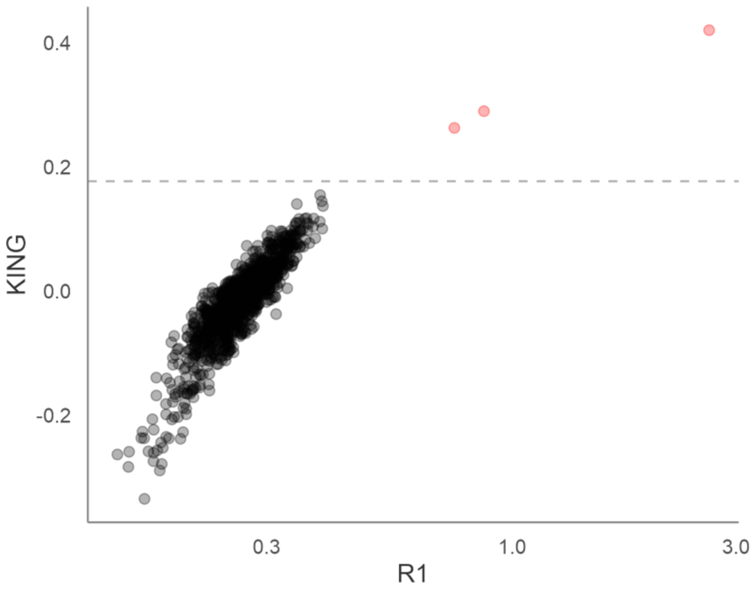


**Figure S1.** Empirical relatedness for all individual pairwise comparisons. R1 coefficients are plotted against KING-robust kinship coefficients. The dashed lines represent the KING-robust kinship and R1 threshold for first degree relatives. Three pairs of individuals fell above this threshold and are colour-coded in red.

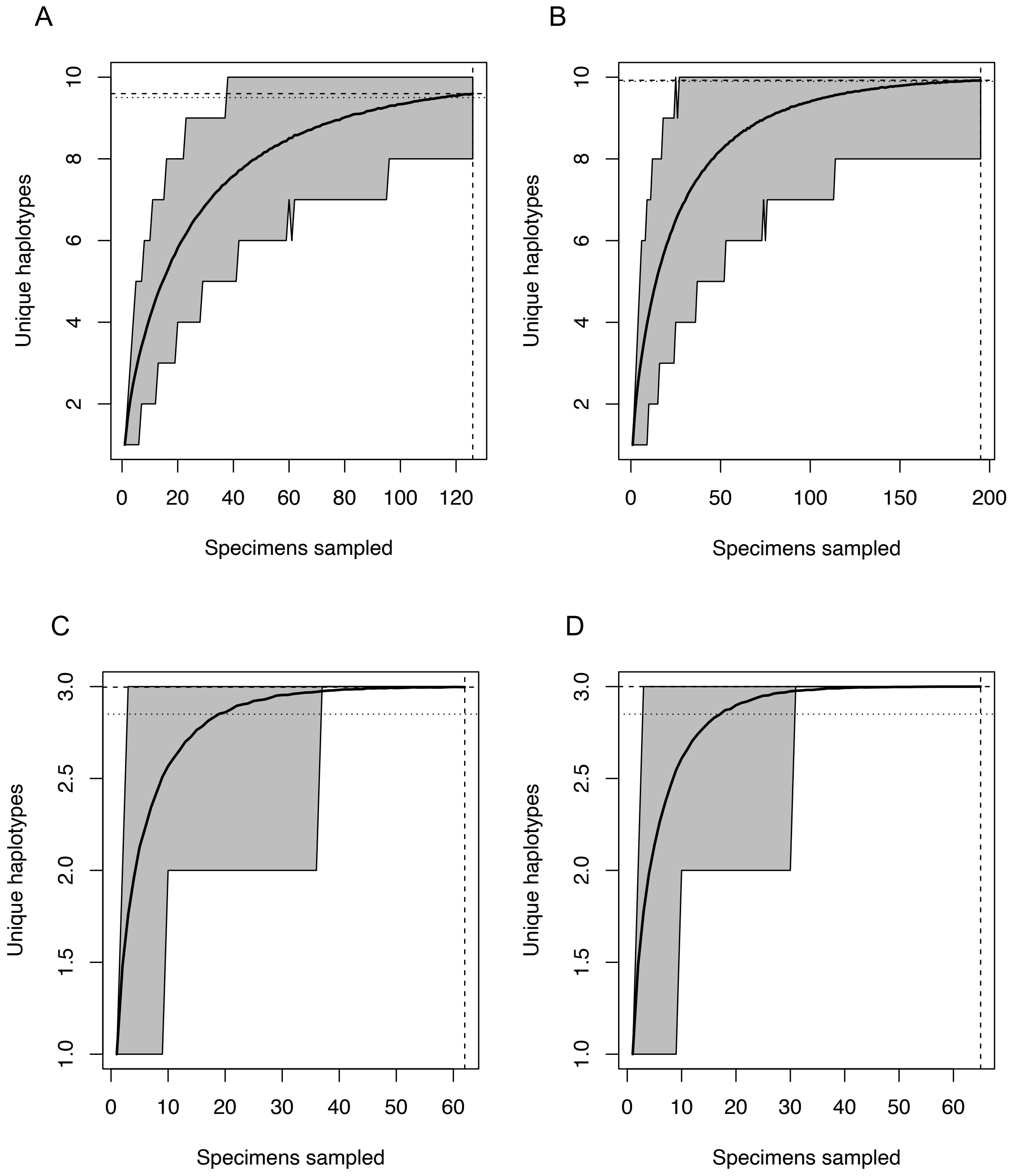


**Figure S2.** Haplotype accumulation curves for (A–B) COI and (C–D) CR sequences identified in *R. ancylostomus.* Grey error bars represent the 95% confidence interval for the number of unique haplotypes. The dashed lines show the observed number of haplotypes and corresponding number of individuals sampled at each iteration of the *HACSim* algorithm. Dotted lines represent the expected number of haplotypes for a haplotype recovery level of (A,C) 95% and (B, D) 99%.

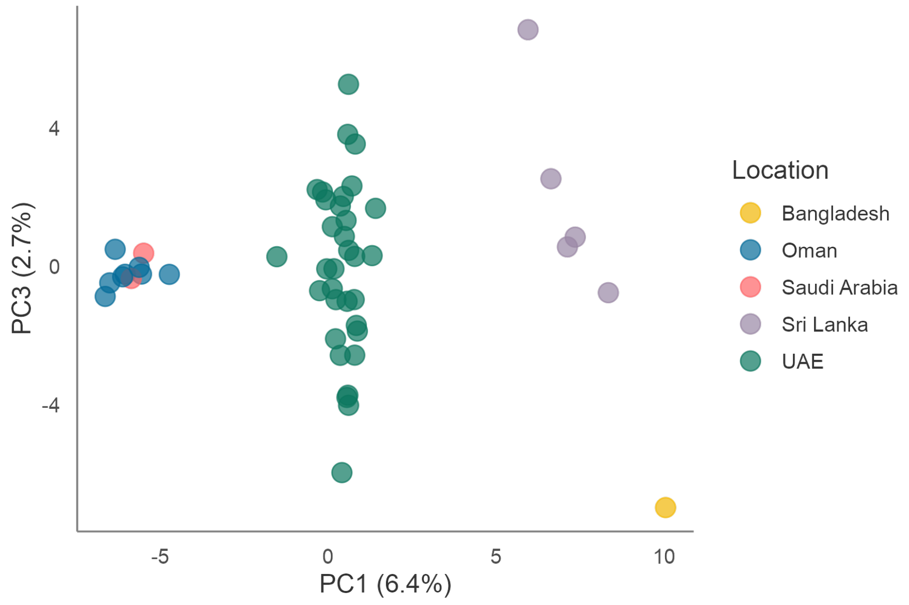


**Figure S3.** Scatterplots showing individual variation in principal components (PC) one and three derived from principal components analysis using 3,565 SNPs and 49 individuals. The amount of variance explained by each PC is shown in parentheses.

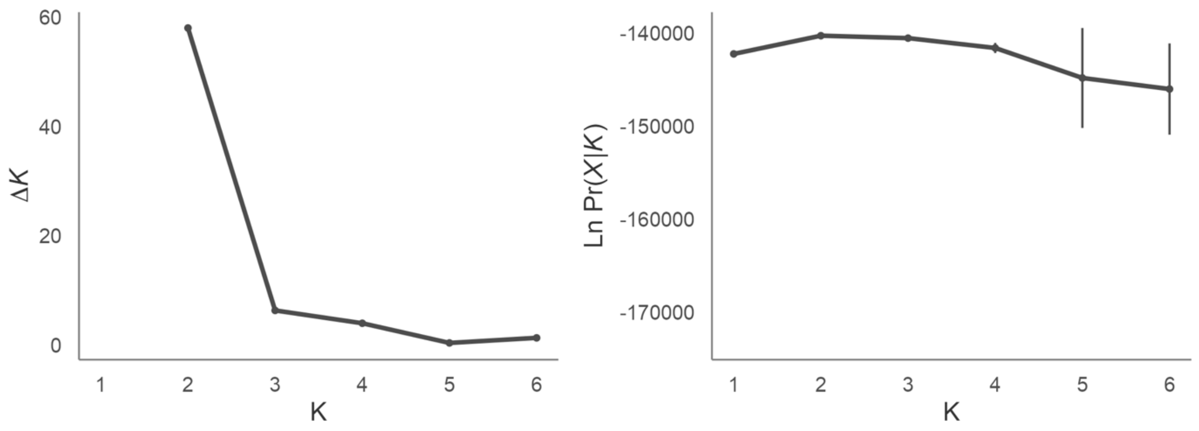


**Figure S4.** (A) Delta K and (B) Ln Pr(*X* | *K*) values with standard errors calculated for 10 replicate runs of STRUCTURE for K = 1 to K = 6 using 3,565 SNPs.

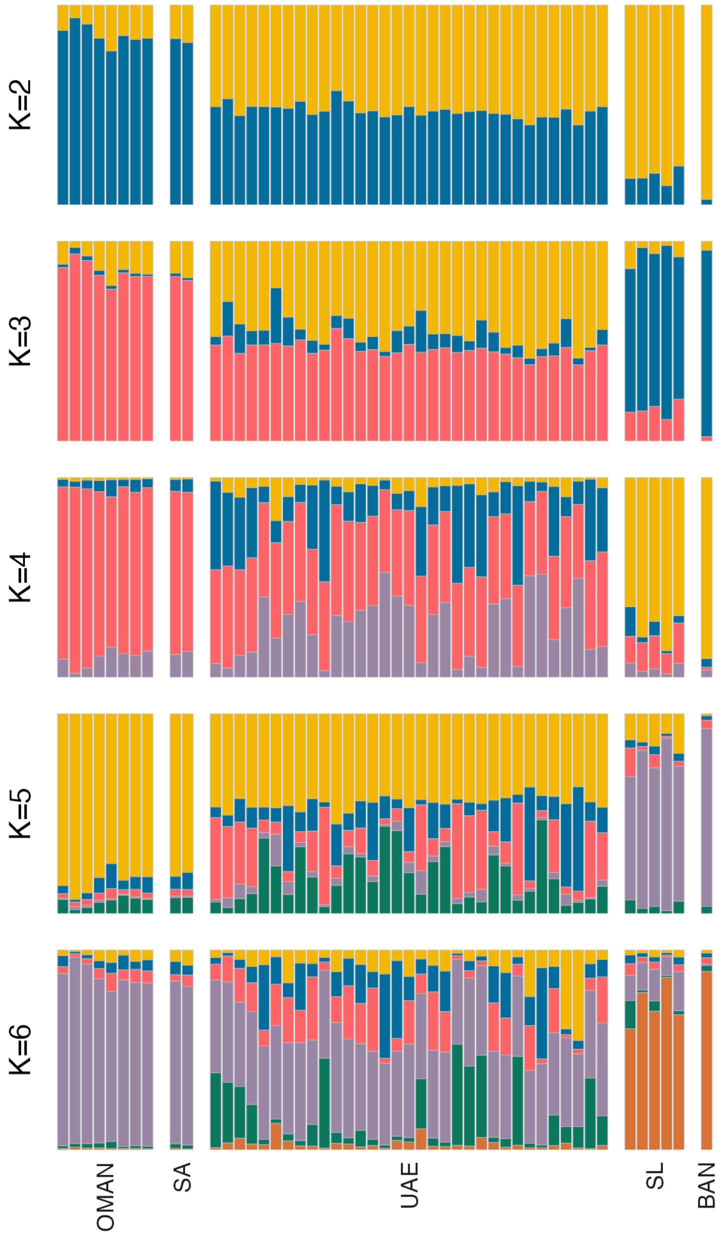


**Figure S5.** Individual assignment to genetic clusters based on STRUCTURE analysis for *K* = 2–6 using 3,565 SNPs and 49 individuals. Each vertical bar represents a different individual and the relative proportion of the different colours indicate the probabilities of belonging to each cluster. Individuals are separated by sampling locations as indicated in Figure 1. Population abbreviations: BAN, Bangladesh; OMAN, Oman; SA, Saudi Arabia; SL, Sri Lanka; UAE, United Arab Emirates.

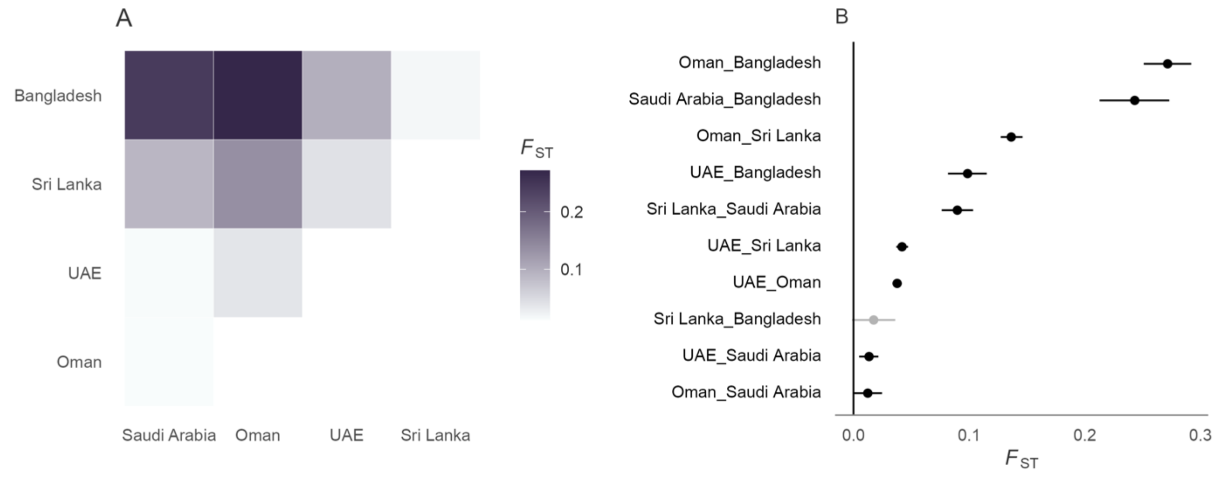


**Figure S6.** (A) Pairwise *F*ST estimates between sampling locations using 3,565 SNPs and 49 individuals. (B) *F*ST estimates and confidence intervals for all pairwise population comparisons. Significant comparisons are indicated in black and nonsignificant comparisons are indicated in grey. *F*ST estimates for population comparisons including Bangladesh are likely to be inflated due to the small sample size of this population.

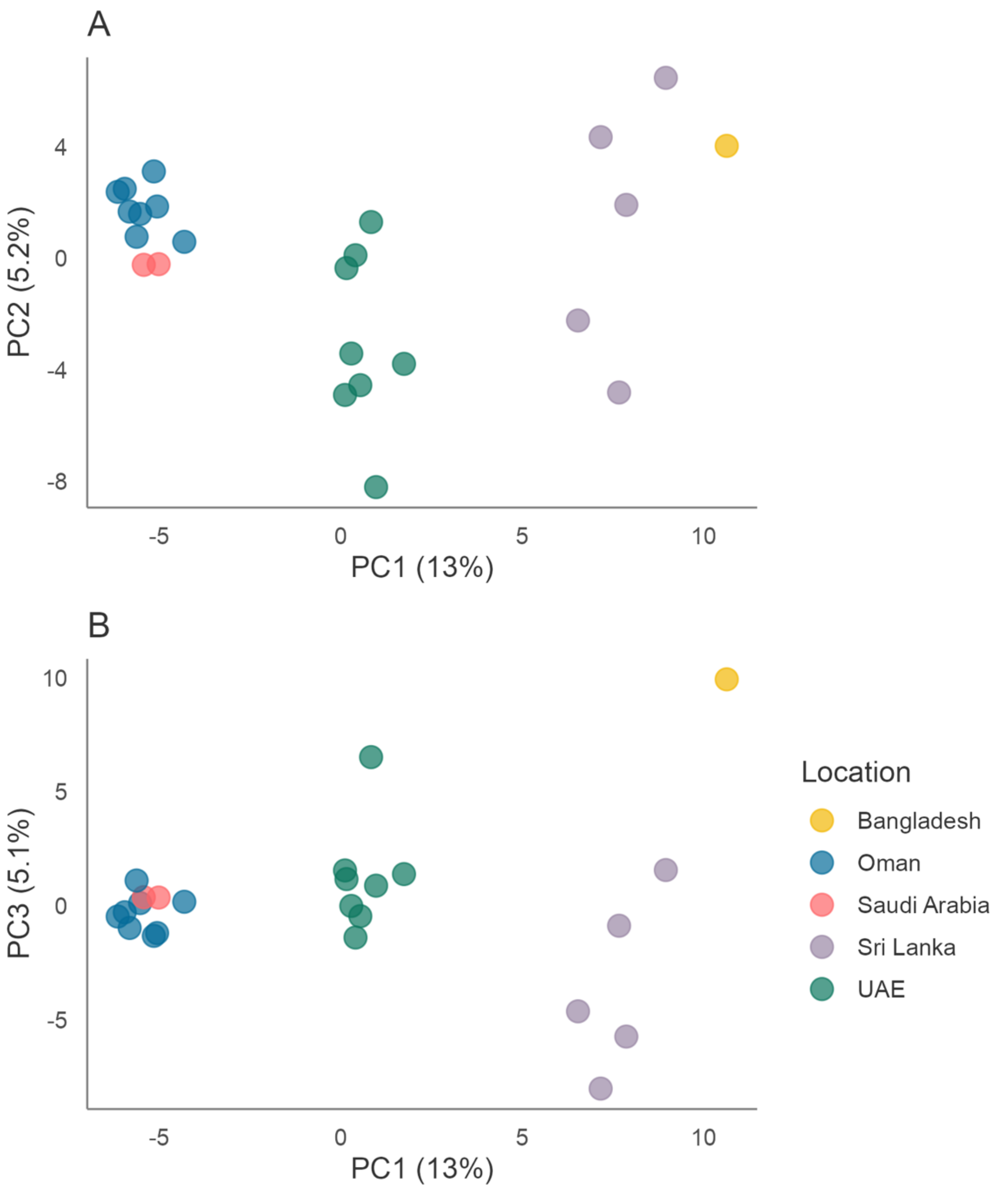


**Figure S7.** SNP-based PCA for a balanced dataset of *R. ancylostomus* where individuals from the UAE were randomly downsampled to $n$ = 8 resulting in a dataset of 3,463 SNPs and 24 individuals. Scatterplots showing individual variation in principal components (PC) one and two (A) and one and three (B) derived from principal components analysis. The amount of variance explained by each PC is shown in parentheses.

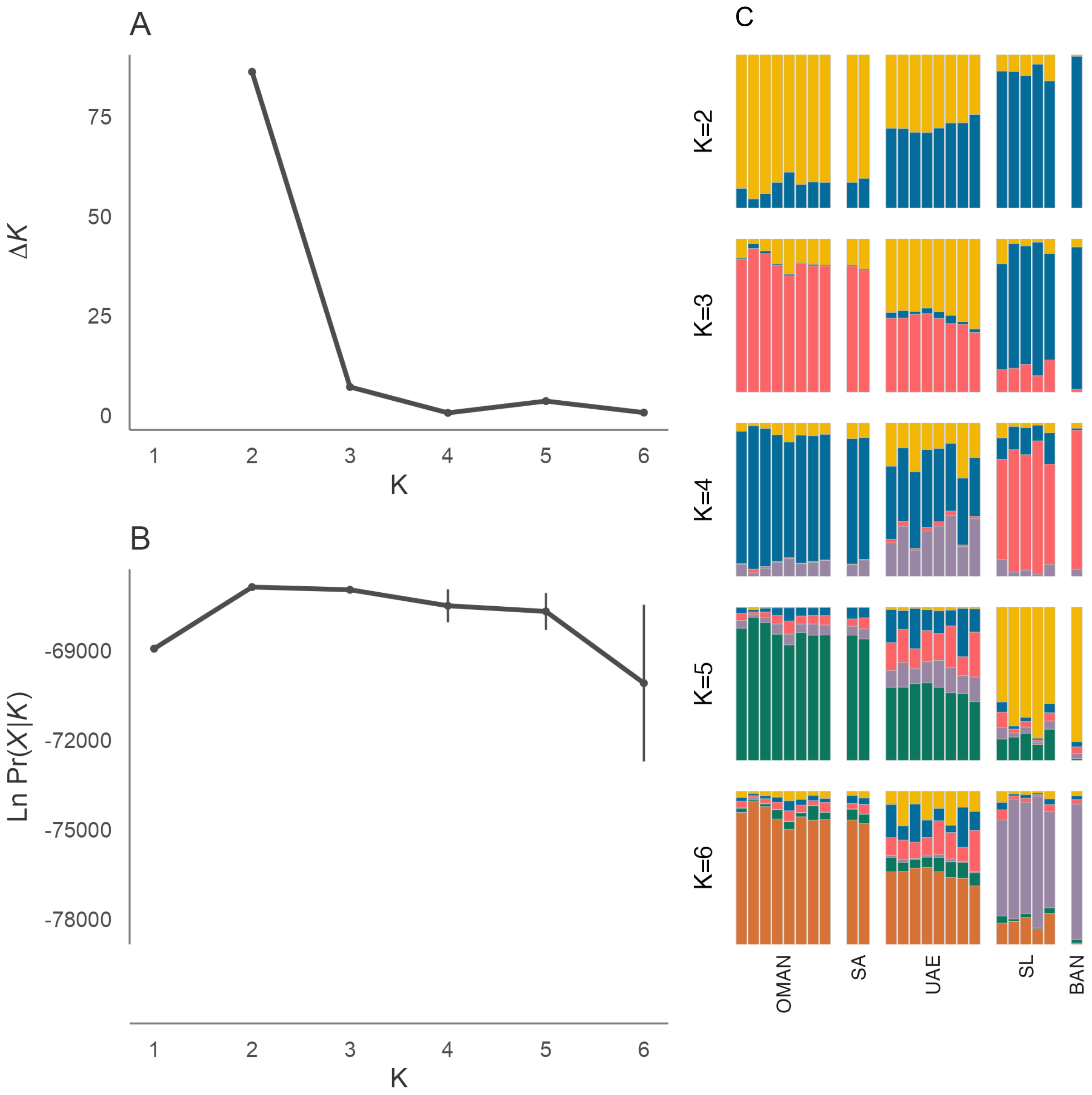


**Figure S8.** SNP-based STRUCTURE analysis for a balanced dataset of *R. ancylostomus* where individuals from the UAE were randomly downsampled to $n = 8$ resulting in a dataset of 3,463 SNPs and 24 individuals. (A) Delta K and (B) Ln Pr($X$ | $K$) values with standard errors calculated for 10 replicate runs of STRUCTURE for K = 1 to K = 6. (C) Individual assignment to genetic clusters based on STRUCTURE analysis for $K$ = 2–6. Each vertical bar represents a different individual and the relative proportion of the different colours indicate the probabilities of belonging to each cluster. Individuals are separated by sampling locations as indicated in Figure 1. Population abbreviations: OMAN, Oman; SA, Saudi Arabia; UAE, United Arab Emirates, SL, Sri Lanka; BAN, Bangladesh.

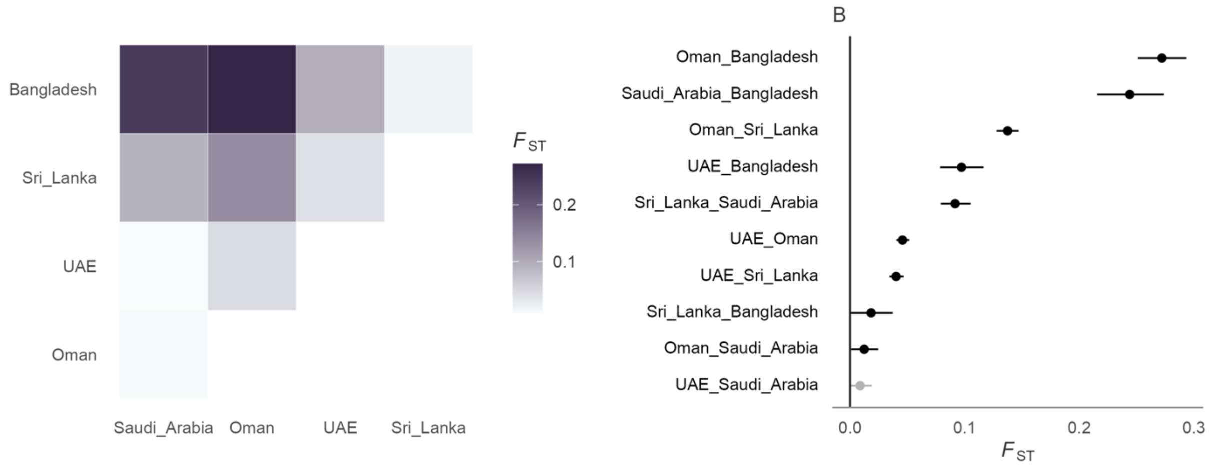


**Figure S9.** (A) Pairwise *F*ST estimates between sampling locations using a balanced dataset of *R. ancylostomus* where individuals from the UAE were randomly downsampled to *n* = 8 resulting in a dataset of 3,463 SNPs and 24 individuals. (B) *F*ST estimates and confidence intervals for all pairwise population comparisons in (A). Significant comparisons are indicated in black and nonsignificant comparisons are indicated in grey. *F*ST estimates for population comparisons including Bangladesh are likely to be inflated due to the small sample size of this population.

### Supplementary Tables

**Table S1**. Sample ID, origin, CR haplotype, COI haplotype and contact email for the samples used in this study. All samples were collected and shipped prior to the species being listed on CITES Appendix II.

| Counter | Sample ID | Origin | COI Haplotype | CR Haplotype | SNP analysis | Contact Email |
|---|---|---|---|---|---|---|
| 1 | 404 | Saudi Arabia | KP_D | KP_B | Y | |
| 2 | 597A | Saudi Arabia | KP_D | KP_B | Y | |
| 3 | 1442 | Saudi Arabia | KP_D | KP_C | Y | |
| 4 | 3000 | Saudi Arabia | KP_D | KP_B | Y | |
| 5 | G173 | Sri Lanka | KP_A | KP_A | Y | |
| 6 | B210 | Sri Lanka | KP_C | KP_B | Y | |
| 7 | B237 | Sri Lanka | KP_D | KP_B | Y | |
| 8 | B242 | Sri Lanka | KP_C | KP_B | Y | |
| 9 | B330 | Sri Lanka | KP_E | KP_A | Y | |
| 10 | G470 | Sri Lanka | KP_C | KP_B | Y | |
| 11 | B564 | Sri Lanka | KP_F | KP_B | Y | |
| 12 | G1384 | Sri Lanka | KP_G | KP_A | Y | |
| 13 | R716 | Sri Lanka | KP_C | KP_B | Y | |
| 14 | 972 | Bangladesh | KP_A | KP_A | N | |
| 15 | Rhina01 | Bangladesh | KP_B | KP_A | Y | |
| 16 | Rhina02 | Bangladesh | KP_B | KP_A | Y | |
| 17 | Rhina03 | Bangladesh | KP_B | KP_A | Y | |
| 18 | RRY164 | Oman | KP_D | KP_B | Y | |
| 19 | RRY222 | Oman | KP_D | KP_C | Y | |
| 20 | RRY223 | Oman | KP_D | KP_C | Y | |
| 21 | RRY225 | Oman | KP_D | KP_B | Y | |
| 22 | RRY859 | Oman | KP_I | KP_B | Y | |

| Counter | Sample ID | Origin | COI Haplotype | CR Haplotype | SNP analysis | Contact Email |
|---|---|---|---|---|---|---|
| 23 | RRY1831 | Oman | KP_I | KP_B | Y | |
| 24 | RRY5872 | Oman | KP_D | KP_B | Y | |
| 25 | RRY5891 | Oman | KP_D | KP_B | Y | |
| 26 | RRY6037 | Oman | KP_D | KP_B | Y | |
| 27 | RRY6149 | Oman | KP_I | KP_B | Y | |
| 28 | RRY6531 | Oman | NA | KP_B | Y | |
| 29 | RRY165 | UAE | KP_D | KP_B | Y | |
| 30 | RRY2039 | UAE | KP_D | KP_C | Y | |
| 31 | RRY2040 | UAE | KP_D | NA | Y | |
| 32 | RRY2182 | UAE | KP_D | KP_B | Y | |
| 33 | RRY2548 | UAE | NA | KP_B | Y | |
| 34 | RRY2980 | UAE | KP_D | KP_B | Y | |
| 35 | RRY3160 | UAE | KP_D | KP_C | Y | |
| 36 | RRY3161 | UAE | KP_D | KP_B | Y | |
| 37 | RRY3162 | UAE | KP_D | KP_C | Y | |
| 38 | RRY3190 | UAE | KP_H | KP_B | Y | |
| 39 | RRY3275 | UAE | KP_D | KP_C | Y | |
| 40 | RRY3572 | UAE | KP_H | KP_B | Y | |
| 41 | RRY3789 | UAE | KP_D | KP_C | Y | |
| 42 | RRY3900 | UAE | KP_D | KP_B | Y | |
| 43 | RRY3901 | UAE | KP_H | KP_B | Y | |
| 44 | RRY4026 | UAE | KP_H | KP_B | Y | |
| 45 | RRY4293 | UAE | KP_J | KP_B | Y | |
| 46 | RRY4884 | UAE | KP_D | KP_C | Y | |
| 47 | RRY4885 | UAE | NA | KP_C | Y | |
| 48 | RRY5219 | UAE | KP_I | KP_B | Y | |
| 49 | RRY6084 | UAE | KP_J | KP_B | Y | |
| 50 | RRY6193 | UAE | KP_D | KP_B | Y | |

| Counter | Sample ID | Origin | COI Haplotype | CR Haplotype | SNP analysis | Contact Email |
|---|---|---|---|---|---|---|
| 51 | RRY6194 | UAE | KP_H | KP_B | Y | |
| 52 | RRY6209 | UAE | KP_D | KP_C | Y | |
| 53 | RRY6249 | UAE | KP_D | KP_B | Y | |
| 54 | RRY6275 | UAE | KP_D | KP_B | Y | |
| 55 | RRY6289 | UAE | KP_D | KP_B | Y | |
| 56 | RRY6290 | UAE | KP_D | KP_B | Y | |
| 57 | RRY6308 | UAE | KP_D | KP_B | Y | |
| 58 | RRY6331 | UAE | KP_D | KP_B | Y | |
| 59 | RRY6348 | UAE | KP_D | KP_C | Y | |
| 60 | RRY6349 | UAE | KP_H | KP_B | Y | |
| 61 | RRY6350 | UAE | KP_D | KP_C | Y | |
| 62 | RRY6429 | UAE | KP_D | KP_B | Y | |
| 63 | RRY6450 | UAE | KP_H | KP_B | Y | |
| 64 | RRY6477 | UAE | KP_H | KP_B | Y | |
| 65 | RRY6478 | UAE | KP_D | KP_C | Y | |
| 66 | RRY6494 | UAE | KP_H | KP_B | Y | |

**Table S2.** Summary showing the number of SNPs and the number of individuals remaining after each step of the SNP filtering pipeline.

| Filter | Number of SNPs remaining | Number of individuals remaining |
|---|---|---|
| Raw genotypes | 19,436 | 65 |
| Low reproducibility scores | 16,376 | 65 |
| One SNP per locus | 15,484 | 65 |
| Read depth < 5 or > 50 | 13,062 | 65 |
| Genotyping rate < 80% | 12,576 | 65 |
| Individual call rate < 96% | 12,576 | 64 |
| Excess heterozygosity | 12,576 | 52 |
| Relatedness | 12,576 | 49 |
| Minor allele frequency < 0.03 | 3,565 | 49 |

**Table S3.** Pairwise $F_{ST}$ estimates (below diagonal) and p-values (above diagonal) for all *R. ancylostomus* population comparisons.

| | Bangladesh | Sri Lanka | Saudi Arabia | UAE | Oman |
|---|---|---|---|---|---|
| | | | COI | | |
| Bangladesh | 0 | 0.01802 | 0.01802 | <0 | <0 |
| Sri Lanka | **0.27407** | 0 | <0 | <0 | <0 |
| Saudi Arabia | **0.75000** | **0.40299** | 0 | 0.28829 | 0.48649 |
| UAE | **0.49757** | **0.32492** | 0.04995 | 0 | 0.09009 |
| Oman | **0.52231** | **0.30033** | 0.07591 | 0.07247 | 0 |
| | | | CR | | |
| Bangladesh | 0 | 0.06306 | 0.01802 | <0 | <0 |
| Sri Lanka | **0.51020** | 0 | 0.46847 | 0.03604 | 0.08108 |
| Saudi Arabia | **0.75000** | 0.00000 | 0 | 0.99099 | 0.99099 |
| UAE | **0.65001** | **0.13640** | -0.15389 | 0 | 0.64865 |
| Oman | **0.76252** | 0.09568 | -0.18736 | -0.02693 | 0 |
